# In humans, insulo-striate structural connectivity is largely biased toward either striosome-like or matrix-like striatal compartments

**DOI:** 10.1101/2024.04.07.588409

**Authors:** AT Funk, AAO Hassan, JL Waugh

**Author notes:** Corresponding author: Jeff Waugh, MD, PhD mailing address: 2222 Medical District Dr., Ste. 105, Dallas, TX, 75235. Authors’ email addresses: AT Funk AAO Hassan.

## Abstract

The insula is an integral component of sensory, motor, limbic, and executive functions, and insular dysfunction is associated with numerous human neuropsychiatric disorders. Insular afferents project widely, but insulo-striate projections are especially numerous. The targets of these insulo-striate projections are organized into tissue compartments, the striosome and matrix. These striatal compartments have distinct embryologic origins, afferent and efferent connectivity, dopamine pharmacology, and susceptibility to injury. Striosome and matrix appear to occupy separate sets of cortico-striato-thalamo-cortical loops, so a bias in insulo-striate projections towards one compartment may also embed an insular subregion in distinct regulatory and functional networks. Compartment-specific mapping of insulo-striate structural connectivity is sparse; the insular subregions are largely unmapped for compartment-specific projections.

In 100 healthy adults, we utilized probabilistic diffusion tractography to map and quantify structural connectivity between 19 structurally-defined insular subregions and each striatal compartment. Insulo-striate streamlines that reached striosome-like and matrix-like voxels were concentrated in distinct insular zones (striosome: rostro- and caudoventral; matrix: caudodorsal) and followed different paths to reach the striatum. Though tractography was generated independently in each hemisphere, the spatial distribution and relative bias of striosome-like and matrix-like streamlines were highly similar in the left and right insula. 16 insular subregions were significantly biased towards one compartment: seven toward striosome-like voxels and nine toward matrix-like voxels. Striosome-favoring bundles had significantly higher streamline density, especially from rostroventral insular subregions. The biases in insulo-striate structural connectivity we identified mirrored the compartment-specific biases identified in prior studies that utilized injected tract tracers, cytoarchitecture, or functional MRI. Segregating insulo-striate structural connectivity through either striosome or matrix may be an anatomic substrate for functional specialization among the insular subregions.

## Introduction

The human insula is a heterogeneous cortical region that is surrounded by the frontoparietal operculum, the temporal operculum, and the Sylvian fissure.^1^ The known functions of the insula are diverse, including interoception, visceral motor, vestibular, heat and pain recognition,^2^ emotional and cardiovascular regulation, and auditory and language processing.^3^ This range of functions is in part mediated by subregional specialization within the insula. The insula can be divided into distinct subregions based on gyral architecture, cytoarchitectural features (neuronal density and subtype), or functional connectivity.^4–8^ For example, cytoarchitectural characterization in multiple mammalian species identified three insular regions distinguished by patterns of neuronal density: the granular (caudodorsal), agranular (rostroventral), and dysgranular (intermediate) subdivisions.^5^ Similarly, segmenting the insula based on covariance in resting state functional connectivity yields three subdivisions: dorsal anterior, with connectivity to frontal, anterior cingulate, and parietal cortices; ventral anterior, with connectivity to limbic and affective areas; and mid-posterior, with connectivity to sensorimotor and nociceptive areas.^6–9^ Insular subregions defined by functional connectivity closely align with segmentations based on cytoarchitectural gradients, underscoring the links between insular structure and function.

The insula displays extensive cortical and subcortical structural connectivity. Insulo-striate connectivity is particularly extensive, is somatotopically organized, and correlates with distinct insular functions, such as reward anticipation and adolescent decision-making.^10,11^ Insulo-striate projections terminate in two structurally and functionally distinct tissue compartments: the striosome and matrix. While both compartments are comprised of medium spiny projection neurons (MSNs), these populations migrate to the striatum at different developmental timepoints^12,13^ via different classes of radial glia,^14^ they have segregated vascular supplies,^15^ they differentially express more than 60 histochemical markers,^16^ and have different afferent and efferent patterns of structural connectivity, as revealed by injected anterograde and retrograde tracers (reviewed in Waugh et al., 2022).

Striosome and matrix appear to be functionally distinct. They are differentially activated during value-driven learning, affective judgements, and action selection.^17–19^ In self-stimulation paradigms, rats acquire and maintain lever-pressing responses significantly more when electrodes contact striosomes than when they contact matrix.^20^ The compartments also have divergent responses to dopamine release^21^ and differential activation following exposure to psychomotor stimulants, such as cocaine and methamphetamine.^16,22–29^ Although the precise location of striosome and matrix varies between individuals, the striosome is generally enriched in the rostro-ventro-medial striatum, while the matrix is concentrated in the caudo-dorso-lateral striatum.^23–28^ Anterograde tracer injections in the rostral and ventral insular cortex of cat and macaque resulted in selective innervation of the striosome, with sparse innervation of the matrix.^25,29^ We recently demonstrated that the human insula follows this same pattern of region-specific compartment bias,^30^ though comprehensive and quantitative mapping of compartment-specific connectivity across all insular subregions was beyond the scope of that publication.

Although the somatotopy of insulo-striate structural connectivity has been studied previously, compartment-specific insular connectivity has not been mapped or quantified, to the best of our knowledge. Given the diversity of functions attributed to the insula, the substantial cytoarchitectural differences between its subregions, and the potential for compartment-specific insulo-striate projections to link neighboring insular subregions with distinct cortico-striato-thalamo-cortical (CSTC) loops,^31,32^ we set out to quantitatively map the structural connectivity of the insular subregions with each striatal compartment. We utilized our previously established diffusion tractography method^30^ in 100 healthy subjects from the Human Connectome Project (HCP) to segment the striatum into voxels with striosome-like and matrix-like patterns of structural connectivity. We then quantified structural connectivity between 19 anatomically segmented insular subregions^33^ and the striosome-like and matrix-like compartments. We found that insulo-striate streamlines that contact striosome-like vs. matrix-like voxels were spatially segregated. The insular zones that displayed compartment-specific bias overlapped with the insular subdivisions previously identified through histologic and MRI-based assessments. This segregation of insulo-striate structural connectivity into compartment-specific networks may contribute to the diverse functions of the human insula, and to the specific symptoms associated with neurodegenerative diseases that affect the insula and/or striatum. Mapping compartment-specific biases in structural connectivity is an essential step in understanding the role of insulo-striate projections in human health and disease.

## Materials and Methods

### 2.1: Overview

In a cohort of 100 healthy subjects, we utilized a previously-established probabilistic diffusion tractography method to identify striatal voxels with striosome-like and matrix-like patterns of structural connectivity.^30^ Following striatal parcellation, we used eight subsequent rounds of probabilistic tractography to validate our striatal parcellations, map compartment-specific insulo-striate streamlines, quantify bias in insulo-striate connectivity, and assess alternate anatomic contributions to the biases in structural connectivity we described. All research was conducted in accordance with the principles set forth in the Declaration of Helsinki and was approved by our University’s Institutional Review Board.

### 2.2: Experimental Cohort

We assembled an experimental cohort of 100 healthy subjects, totaling 200 hemispheres, from the Human Connectome Project dataset,^34–36^ accessed through the National Institute of Mental Health Data Archive (NDA).^37^ This cohort was racially diverse, included broad representation across the adult age span, and was balanced for sex. We identified ten HCP subjects (five female, five male) at five-year intervals, beginning at 20 years old and ending at 65 years old (20, 25, 30, 35, etc.). If a particular age block had an insufficient number of HCP subjects to reach our 10-subject goal, we searched for participants in adjacent ages, but never crossed into an adjacent five-year interval. Within each 10-subject block (5 female, 5 male), we aimed to create a diverse racial representation by including at least one Asian and at least one Black individual (as defined by the subject) for each sex at each age block. For any age-sex block in which an insufficient number of subjects who identified as a particular race was available, we overfilled either the adjacent age block or the opposite sex from the same age block. We filled the remainder of each age/sex block with White subjects, which were abundant in the HCP database. In 6 of 10 age blocks, there were insufficient subjects available to meet all of our target criteria (a diverse group of 5 females, 5 males), so we overfilled the opposite sex for that age block to reach the 10-subject goal and balanced such mismatches in adjacent age blocks. This 100-subject cohort was therefore evenly split between females and males (50:50). Subjects self-reported their race as: 13% Asian, 28% Black, 51% White, and 8% Multiracial/Other/Not reported. The average age of these subjects was 42.3 years (age range: 20-65 years). Based on the Edinburgh Handedness Inventory,^38^ 90 subjects were right-handed (90%) and 10 were left-handed (10%). All 100 subjects were healthy, with no recognized neurologic or psychiatric conditions. We imposed a pre-set criteria for including a hemisphere in our final analyses (see 2.4: Insular Segmentation, below), which eliminated four hemispheres. No subject had both hemispheres eliminated.

### 2.3: Background - Striatal Parcellation Method

In 2022, we described a method to parcellate the striatum into striosome-like and matrix-like compartments *in vivo* in humans.^30^ We further developed this method to assess compartment-specific striato-pallido-thalamic projections, demonstrating that most thalamic nuclei have compartment-specific biases in structural connectivity.^39^ In brief, striatal parcellation leverages the anatomic findings from four decades of histologic investigation in animals, which used injected tract tracers to map compartment-selective projections in mice, rats, cats, dogs, and non-human primates. We selected extra-striate regions with compartment-selective afferents in animals and organized them into groups that favored either the striosome or matrix compartments. We used these groups as “bait” regions for quantitative probabilistic tractography. For each striatal seed voxel, the abundance of streamlines that reached striosome-favoring vs. matrix-favoring bait regions defined that voxel’s compartment-specific connectivity profile. We selected the most-biased voxels (those whose connectivity bias was ≥1.5 standard deviations above the mean for the striosome-like and matrix-like probability distributions) to represent the compartments in subsequent quantitative assessments of insulo-striate structural connectivity. Given the inferential nature of this method, we henceforth refer to parcellated striatal voxels as “striosome-like” or “matrix-like.”

The reliability of connectivity-based parcellations was high; test-retest comparisons in scans performed one month apart identified an error rate of 0.14%.^30^ Voxels identified as striosome-like or matrix-like were unlikely to convert between compartments in a subsequent scan. We defined these voxels based solely on their connectivity profiles, but then assessed other anatomic features of the striosome and matrix (as reported from the animal and human histology studies cited above) to check the validity of these connectivity-based striatal parcellations: their spatial distribution, relative abundance, and connectivity with regions that were mapped in animals but were left out during striatal parcellation. In every case, our striosome-like and matrix-like voxels shared the anatomic features demonstrated through injected tract tracers and histochemistry. However, we urge readers to bear in mind that this method is probabilistic and indirect – we have not directly identified striosome and matrix tissue.

### 2.4: MRI Acquisition and Processing

This study was a secondary analysis of existing HCP MRI data. All subjects were scanned at 3 Tesla with a whole-brain diffusion tensor imaging (DTI) protocol. All MRI data was collected in a single scan session. All subjects were scanned using Siemens Prisma scanners running Syngo MR E11 software and using harmonized protocols at three separate sites in the United States.^35^ DTI for these subjects was obtained at 1.5 mm isotropic resolution using 200 diffusion directions (14 B_0_ volumes, 186 volumes at non-colinear directions) with the following parameters: repetition time = 3.23 seconds; echo time = 0.0892; imaging volume = 256×256×256 mm. We utilized paired anterior-posterior and posterior-anterior DTI volumes to estimate and correct for susceptibility-induced field distortions using the FSL tool *topup*. We performed skull stripping and motion correction using the FSL Brain Extraction Tool (*bet2*). We utilized the FSL tool *eddy_openmp* to correct for eddy current distortions and head motion. We then used the FSL tool *bedpostx* to fit probability distributions on diffusion parameters at each voxel, including modeling of crossing fibers. Using *dtifit*, we fit local diffusion tensors to create 3D FA images at 1.5 mm isotropic resolution. We completed all DTI preprocessing and tractography steps in each subject’s native diffusion space, utilizing our University’s distributed Linux cluster.

### 2.5: Insular Segmentations

We utilized the insular segmentations of Ghaziri et al.^33^ to extract connectivity measures from probabilistic tractography. We selected this atlas because the ROI segmentation offered substantial granularity, its subregions conformed to human gyral anatomy, and it was derived using the same (diffusion) methodologies utilized in the present study. This segmentation technique parcellated the human insular cortex into 19 distinct subregions for each hemisphere using a K-means random parcellation algorithm followed by manual segmentation of the insula to harmonize subregions with sulco-gyral divisions. These segmentations were graciously shared by the D. Nguyen lab, University of Montreal, Canada. We used these subregional segmentations in MNI standard space (1 mm isotropic voxels) to extract connectivity measures. We determined that registration of these insular subregion masks into native diffusion space led to boundary erosion between the adjacent masks, leading to occasional gaps or overlap. If we had measured insulo-striate connectivity in native space, this loss of precision would yield incomplete and inaccurate sampling. Therefore, we registered tractography-derived probability maps (that were generated in native diffusion space) into MNI standard space for quantification of tractography for each insular subregions.

### 2.6: Anatomic Masks

We defined seed, waypoint, and bounding masks in MNI standard space and subsequently registered these masks into subjects’ native diffusion space. All tractography was completed in native diffusion space. We manually segmented the caudate and putamen, excluding the nucleus accumbens. We also excluded the caudal half of the tail of the caudate, as we found that registration errors and partial volume effects reduced the accuracy of registration for this narrow tail.^30^ We generated a whole-insula seed for tractography by summing all insular subregions,^33^ inflating that volume by one voxel in all dimensions, and then manually trimmed the mask relative to the MNI152_T1_1mm standard. We generated an insula-subcortical bounding mask to refine our tractography, which encompassed the insula, caudate, putamen, and the white matter surrounding these structures. We utilized pilot rounds of tractography to refine this bounding mask, and visually inspected each subject and hemisphere to ensure that this mask did not exclude valid insulo-striate streamlines. This mask eliminated all streamlines that extended beyond its boundaries, excluding non-insular corticostriate, striatopallidal, thalamostriate, and striatal-brainstem projections. All other regions of interest (ROIs), including bait regions, striatal segmentations, and our midline exclusion mask, were identical to those detailed in our original description of this method ^30^. The bait, seed, waypoint, and bounding masks can be accessed here: github.com/jeff-waugh/Striatal-Connectivity-based-Parcellation.

### 2.7: Probabilistic Tractography

We completed seven rounds of probabilistic tractography in this study (Table 1). We will briefly summarize the goals of each round, then provide a deeper background on each round of tractography in subsequent paragraphs. First, we parcellated the striatum into striosome-like and matrix-like voxels, based on differential structural connectivity (tractography round 1, see Methods 2.3). We measured anatomic features (detailed below) of these parcellated striatal voxels to assure that our striosome-like and matrix-like masks shared the anatomic features of striosome and matrix measured in tissue. Tractography rounds two and three were part of this validation step. We assessed whether compartment-level biases in cortico-striate projections matched the biases measured through injected tract tracers in animals: parcellating with one bait region left out (an N-1 parcellation, tractography round two), followed by quantification of compartment-specific bias in the left-out region (round three). These voxels served as the seeds or targets for rounds of tractography that tested our hypotheses (rounds 4-7). Round four mapped the location of insulo-striate streamlines, comparing streamlines that targeted striosome-like vs. matrix-like targets. Round five quantified compartment-specific biases in insulo-striate structural connectivity for each of the 19 insular subregions. Rounds 6-7 were post-hoc tests of alternative hypotheses, assessing non-compartment drivers of connectivity bias. Round six considered the possibility that nucleus-of-origin (caudate vs. putamen) might influence compartment-specific biases in insulo-striate connectivity. Round seven tested the hypothesis that local anatomic features of the striatum, separate from compartment-level bias, might explain our insulo-striate findings.

**Table 1:** Summary of the seven rounds of tractography utilized to test our experimental aims. Rounds were organized sequentially, based on the experimental goals that served those aims. “Type of Tractography” refers to either traditional streamline tractography or classification targets tractography. Rounds 1-3 (green) established and validated our striatal compartment parcellations, rounds 4 and 5 tested our primary hypotheses regarding insulo-striate connectivity (white), and rounds 6 and 7 tested potential alternate explanations for our findings (blue). Abbreviations: CTT, classification targets tractography; Caud, caudate; Put, putamen.

| Round | Type of Tractography | Seed Region | Target Region | Goal of Tractography |
| --- | --- | --- | --- | --- |
| 1 | CTT | Striatum | Extra-striate Bait Regions | Parcellate striatum into striosome-like, matrix-like voxels |
| 2 | CTT | Striatum | Extra-striate N-1 Bait Regions | Test accuracy of striatal parcellations from Round 1 |
| 3 | CTT | N-1 Region | Compartment-specific N-1 Striatal Masks | Test accuracy of striatal parcellations from Round 1 |
| 4 | Streamline | Whole Insula | Compartment-specific Striatal Masks | Map compartment-specific insulo-striate streamlines |
| 5 | CTT | Whole Insula | Compartment-specific Striatal Masks | Quantify biases in insulo-striate connectivity |
| 6 | CTT | Whole Insula | Caud/Put-Specific Compartment Masks | Assess compartment-specific vs. nucleus-of-origin effects |
| 7 | CTT | Whole Insula | Compartment-adjacent Striatal Masks | Assess compartment-specific vs. neighborhood effects |

All rounds of tractography were executed in each subject’s native diffusion space. We utilized the following probtrackx2 settings for all tractography: curvature threshold = 0.2; steplength = 0.5 mm; number of steps per sample = 2,000; distance correction, to prevent target proximity from influencing connection strength. Rounds one and two (striatal parcellation) used a whole-hemisphere mask to bound the tractography; rounds three through nine (insulo-striate tractography) used the previously described insulo-striate bounding mask that restricted streamlines to only those transiting the insula, striatum, and surrounding white matter, eliminating streamlines that strayed outside the insulo-striate path. Rounds one, two, and four produced adequate samples by seeding with 5,000 streamlines per seed voxel, while rounds three, and 5-7, required an increase to 50,000 streamlines per seed voxel to adequately quantify connectivity within their seed volumes.

In tractography round one, we used classification targets tractography (CTT) to parcellate the striatum into striosome-like and matrix-like voxels (see Methods 2.3). Briefly, we used sets of striosome-favoring and matrix-favoring regions as “bait” for CTT. We previously demonstrated that this set of bait regions, which included the rostral insula, produced robust striatal parcellations.^30^ However, one cannot use a region to parcellate the striatum and then accurately quantify connectivity with that region. We therefore removed the insula from the set of bait regions and performed striatal parcellation with the remaining four striosome-favoring regions (mediodorsal thalamus, posterior orbitofrontal, basolateral amygdala, and basal operculum) and five matrix-favoring regions (supplementary motor area, primary motor cortex, primary sensory, globus pallidus interna, and the VLc/VPLo thalamic nuclei). This parcellation method is identical to one that we utilized previously,^30^ identified there as an “N-1 analysis.”

Following striatal parcellation, we selected voxels from the tails of the probability distribution to generate equal volume masks of the most-biased striosome-like and matrix-like voxels. To eliminate the possibility that differing target size could skew streamline quantification, we ensured that the striosome-like and matrix-like masks in each hemisphere, and in each individual, were always of equal volume. We set a fixed volume standard for inclusion (the uppermost 13% of voxels from each distribution, equivalent to 1.5 standard deviations above the mean in a normal distribution) from each subject’s striosome-like and matrix-like probability distributions. These striosome-like and matrix-like masks were utilized, in subsequent steps, as the seeds or targets for other rounds of tractography.

Before proceeding to insulo-striate mapping, we compared the features of our striatal parcellations with the expected relative abundance and location of striosome and matrix, established previously through histology. We measured the volume of striosome-like and matrix-like voxels that reached substantial bias (connection probability ≥0.87, to parallel the 1.5 standard deviation threshold noted above). For each hemisphere and subject, we expressed these measures as the percentage of all supra-threshold volume (compartment X / (compartment X + compartment Y)). Since the distributions of striosome and matrix differ in their intrastriate location (as demonstrated through histology), we measured location of these compartment-like voxels as another way to assess the accuracy of our parcellations. For each voxel in our striosome-like and matrix-like masks, we measured its cartesian position relative to the centroid of the nucleus it resided in (caudate or putamen) for each subject and hemisphere, producing a dataset of 66,812 parcellated voxels. Finally, we quantified compartment-specific bias in structural connectivity for primary motor cortex and posterior orbitofrontal cortex, regions with robust matrix-favoring and striosome-favoring bias, respectively.^24,25,40^ Just as we performed N-1 parcellations to leave out the rostral insula (tractography round one), for tractography round two we performed striatal parcellation with either the primary motor or posterior orbitofrontal cortices left out. We then identified the most-biased voxels for each of these N-1 parcellations (equal volume striosome-like and matrix-like masks, distinct from those generated for tractography round one) to serve as targets for quantitative CTT (tractography round three).

Tractography round four mapped insulo-striate streamlines with two iterations, alternating the seed and target ROIs (A-to-B and B-to-A) and averaging the results (AB-BA). First, we used the whole insula as seed and either striosome-like or matrix-like voxels (in separate iterations) as an obligatory waypoint. Second, we set striosome-like or matrix-like voxels as seeds (in separate iterations) and the whole insula as an obligatory waypoint. We performed this round of tractography in both directions (whole insula to compartment-specific voxels, and compartment-specific voxels to whole insula) to reduce potential bias from the directions of diffusion acquisitions and to increase the reliability of streamline mapping. We then averaged the output of these two iterations (A-to-B–B-to-A) for each subject and hemisphere. The output of this round of tractography (streamline distributions that were largely extra-insular) allowed us to quantify and map streamline bundles that connected the insula to each striatal compartment. The striosome-like and matrix-like versions of these runs of tractography utilized the same insular seed mask, seeded the same number of streamlines per insular voxel, were bounded by the same insulo-striate bounding mask, utilized equal volume striatal masks, and corrected for any path length differences between seed and target voxels.

In tractography round five, we mapped compartment-specific connectivity within the insula by performing CTT with the insula as seed and the two striatal compartments as targets. The output of this round of tractography, striosome-favoring and matrix-favoring probability maps of the insula, allowed us to quantify connectivity at each insular voxel, and thus for each insular subregion. We noted that utilizing the standard *probtrackx2* settings for streamline seeding (5,000 streamlines per seed voxel) left some subjects, and some subregions of the insula, relatively undersampled. Increasing the depth of sampling to 50,000 streamlines/voxel yielded a more robust probability distribution, and therefore, more accurate quantification of connectivity. Specifically, we noted in subsequent quantification steps that artificial binarization of the connectivity data (when suprathreshold voxels were all striosome-like, or all matrix-like) led to less accurate connectivity estimates. Increasing the number of streamlines per seed voxel reduced the number of subjects and insular subregions with zero connectivity for one striatal compartment, and thus reduced this artificial binarization.

After assessing our primary aims with these five rounds of tractography, we recognized that our measures of compartment-specific insular connectivity might have been influenced by relative differences in regional connectivity – that striosome vs. matrix differences might have been influenced by caudate vs. putamen differences in structural connectivity. We therefore carried out a post-hoc round of tractography (round six) aimed at understanding the influence of caudate/putamen connectivity on striosome/matrix connectivity. Round six differed from round five only in the location of the compartment-like target masks. For this iteration, we used the same striatal parcellations (probability maps) we generated in tractography round one but defined new striosome-like and matrix-like masks. Rather than choosing the most-biased voxels (uppermost 1.5SD) from the whole striatum, we defined these masks proportionally, adjusting for the relative volume of caudate and putamen. Specifically, instead of choosing the 180 most-biased voxels from any part of the striatum, we chose the 80 most-biased voxels from the caudate and 100 most-biased voxels from the putamen. We assured that within every combination of subject, hemisphere, and region, the volume of striosome-like and matrix-like voxels was equal. We utilized these regionally proportionate compartment masks as targets for CTT, allowing us to map connectivity between each insular voxel and the striatal compartments without the potential for caudate-putamen bias.

In tractography round seven, our goal was to distinguish compartment-specific effects from the influence of the striatal “neighborhood” in which those voxels reside. That is, we wished to learn whether other voxels that were similarly positioned, but that were not selected based on biased connectivity, would also drive the biases in insulo-striate connectivity seen in tractography round five. Round seven was identical to round five – CTT with the whole insula as seed – with one exception: instead of targeting the precisely-selected striosome-like and matrix-like masks, we targeted their near-neighbors. For every voxel in the compartment-like masks, we shifted the position in each plane by ±0-3 voxels at random, ensuring that no voxels in the randomly shifted mask were reselected from either original 1.5SD mask. This randomized relocation ensured that the precise-to-neighboring voxel change would maintain the topographic organization of our compartment-specific 1.5SD masks. We measured the position of every shifted voxel for comparison with the original 1.5SD voxels to assure that our masks had not shifted, on average, from the original “neighborhood.” We then performed quantitative CTT with the whole insula as seed and these location-shifted neighboring striatal voxels as targets.

### 2.8: Assessing the Amplitude and Location of Insulo-striate Streamlines

As we utilized two different modes of tractography (traditional streamline in round four vs. CTT in rounds 1-3 and 5-9) we required two different approaches to quantify the results of these rounds of tractography. For tractography round four, we quantified the total number of insula-seeded streamlines that reached striosome-like or matrix-like voxels. We then normalized that count by the volume of that individual’s insular seed mask to reduce the impact of inter-individual differences in insula size. We compared this count – the number of streamlines per insular voxel – between striosome-favoring and matrix-favoring tractography for left and right hemispheres. The relative abundance of striosome-favoring vs. matrix-favoring streamlines was biased toward the same compartment in both hemispheres, so we combined the hemispheres for subsequent quantification steps. Since we were investigating biases in connectivity, rather than the absolute strength of connectivity, we expressed connectivity as the percentage of streamlines that reached matrix-like voxels (matrix count/(striosome count + matrix count)). This formulation reduced the outsize influence of subjects whose total streamline counts were unusually high or low.

After registering each subject’s tractography into MNI space, we generated averaged streamline tractography volumes for each hemisphere and compartment. Distinguishing between the tract core and the penumbra of less-specific streamlines is critical when assessing the streamlines of probabilistic tractography ^41,42^ ^43^ ^44^. We therefore normalized each averaged tract relative to its maximum amplitude and set thresholds to retain the uppermost 25%, 50%, or 75% of all voxels. We then assessed the volume of each thresholded average, and their volume of overlap, to determine the Dice similarity coefficient (DSC), as follows:

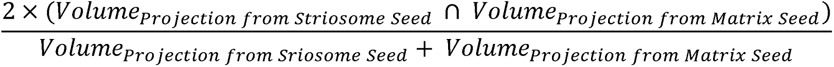

### 2.9: Assessing Compartment-specific Bias in Insular Subregions

For tractography iteration five (CTT, insula as seed, striosome-like and matrix-like voxels as targets), we assessed the intra-insular locations of compartment-specific connectivity, as well as the mean connectivity bias for each of 19 insular subregions. First, we visually inspected the averaged insular probability distributions for striosome-favoring and matrix-favoring CTT. While matrix-favoring voxels occupied a single cluster, striosome-favoring voxels occupied two, separated clusters in the rostral and caudal insula. We chose to assess these two clusters of striosome-favoring voxels independently, with each cluster compared to the position of the matrix-favoring cluster. For each individual and hemisphere, we identified the center of gravity (COG, using *fslstats*) within the matrix-favoring distribution, spanning the whole insula. For the striosome-favoring distribution we split the insula at coronal plane y=1 (MNI convention), the center of the matrix cluster and therefore the nadir of the striosome-favoring distribution. We then identified the COG within the rostral and caudal striosome-favoring distributions for each individual and hemisphere. We adjusted the cartesian position of all COG measures to account for differences in the position of the insula between left and right hemispheres, so that we could combine left and right hemisphere data when measuring inter-compartment differences in COG location.

### 2.10: Determining Compartment Specificity for Insular Subregions

Next, we assessed the degree of compartment specificity in each of 19 insular subregions. To improve the accuracy of our data extraction, we first corrected for partial volume effects induced by registration of probability maps into MNI space. In native space, the striosome-like and matrix-like probability distributions sum to one at each voxel. After registration to MNI space, probability at some voxels no longer summed to one. Therefore, we renormalized the striosome-favoring and matrix-favoring probability distributions on a voxel-by-voxel basis. We trimmed edge voxels whose summed value (striosome plus matrix) was <0.5, which reduced partial volume effects at the edges\ of the insula. After normalization and edge trimming, we utilized the insular segmentations of Ghaziri et al. (2018) to extract data for each of 19 insular subregions. We quantified the number of suprathreshold voxels (value ≥0.55) in striosome-favoring and matrix-favoring normalized probability distributions within each insular subregion. For each subregion, we expressed connectivity biases as the volume percent: the percent of suprathreshold voxels dominated by each compartment (N_voxels_, striosome or matrix / (N_voxels_, striosome + N_voxels_, matrix). For all insular subregions, the left and right hemispheres were biased toward the same compartment, so we combined hemispheres for this quantification step.

Our method of quantifying compartment-specific bias could be compromised by undersampling of the probability distribution. For example, consider the addition of one marginal voxel in two subjects with the same compartment ratio – subject A, with 1 matrix-favoring voxel and 9 striosome-favoring voxels, and subject B, with 10 matrix-favoring voxels and 90 striosome-favoring voxels. Adding a single matrix-favoring voxel to each subject will increase the matrix bias by 8.2% for subject A, and 2.1% for subject B. To avoid non-linear influences on quantification, we set a minimum threshold of 19 suprathreshold voxels (the number of insular subregions considered) for each compartment-specific probability distribution. That is, to be included in our final cohort for quantification a hemisphere had to have at least 19 suprathreshold voxels for both striosome-favoring and matrix-favoring distributions. This minimum sample criteria led us to eliminate four hemispheres, for a final sample of 196 hemispheres. Note that the increase in the number of streamlines per seed voxel to 50,000 (a 10x increase over the standard settings) substantially decreased the number of undersampled hemispheres.

Tractography rounds six and seven were post-hoc analyses to determine the influence of regional connectivity biases (caudate vs. putamen; precisely selected vs. neighborhood voxels) on compartment-specific connectivity biases. For these rounds, we performed tractography with the same parameters as round five (CTT with insular seed and striatal compartment targets), but with seed or targets adjusted to test alternate explanations for our insulo-striate findings. As in round five, for rounds six and seven we expressed compartment-specific connectivity as the volume percent projecting to either striosome-like or matrix-like voxels.

### 2.11: Significance Testing

We performed significance testing using STATA (StataCorp, 2023, Stata Statistical Software: Release 18. College Station, TX). We completed two sets of comparisons to assess the accuracy of our striatal parcellations: intrastriate voxel location, and compartment-specific bias in cortico-striate structural connectivity (N-1 analyses). We used two-factor ANOVA to assess the effect of striatal compartment and nucleus of origin (caudate or putamen) on intra-striate voxel location (tractography round one; separate ANOVAs for the x-, y-, and z-planes). Since subjects’ compartment-like mask volume differed from person to person depending on the availability of highly biased, compartment-specific voxels, we controlled for subject identity as a nuisance variable. We previously demonstrated that scanner type, subject sex, and self-identified race had no significant influence on voxel location.^30^ Therefore, we did not model these factors. We did not include hemisphere as a factor, as no interhemispheric differences in matrix and striosome location have been described previously, to the best of our knowledge. We analyzed simple main effects for factor interactions using the SME utility (UCLA ATS Statistical Consulting Group).^45^ We used the conservative simultaneous test procedure for estimating F-critical available through the SME utility. We assessed compartment-specific biases in cortico-striate connectivity (tractography round three) using two t-tests. We compared striosome-favoring and matrix-favoring voxels (volume percent) for two regions, primary motor and posterior orbitofrontal cortices, and therefore utilized a significance threshold of p = 0.025 (Bonferroni correction for these and all subsequent comparisons; 0.05 / 2 tests).

We compared insulo-striate streamline counts between the hemispheres (tractography round four) using three t-tests (two tailed, paired samples; left vs. right hemisphere, total streamlines; left vs. right hemisphere, matrix-favoring streamlines; left vs. right hemisphere, striosome-favoring streamlines). We therefore set a significance threshold of p = 0.0167 (0.05 / 3 tests). We compared the location and amplitude of streamline bundles (those that favored striosome-like vs. matrix-like voxels) using voxelwise, non-parametric testing (FSL’s *randomise*) with the following parameters: 5,000 permutations; variance smoothing = 2 mm; threshold-free cluster enhancement mode; masked by the same subcortical bounding mask utilized to generate the streamlines (tractography round four). For each subject we combined left and right hemisphere tractography into a single bilateral volume to reduce the number of *randomise* comparisons. As we completed a subsequent round of *randomise* testing with similar goals (described below), we considered these a family of tests and therefore utilized a significance threshold of p = 0.025 (0.05 / 2 tests). For all *randomise* comparisons, we utilized familywise error-correction and threshold-free cluster enhancement.

We compared the intra-insular root-mean-square distance between centers of gravity for striosome-favoring and matrix-favoring clusters (tractography round five) using one sample t-tests with the assumption of no location difference between compartment-specific centers of gravity. We carried out two of these one-sample tests (anterior striosome-favoring vs. matrix-favoring; posterior striosome-favoring vs. matrix-favoring), so we set our significance threshold for root-mean-square comparisons at p = 0.025 (0.05 / 2 tests). We compared the x-, y-, and z-plane locations for the COGs of striosome-favoring and matrix-favoring clusters, testing anterior striosome-favoring vs. matrix-favoring, and posterior striosome-favoring vs. matrix-favoring COGs. We therefore carried out six t-tests (two tailed, paired samples), so set our significance threshold at p = 8.3 × 10^−3^ (0.05 / 6 tests).

We tested each insular subregion for compartment-level biases in structural connectivity through two related, but distinct approaches. Both approaches utilized the classification targets probability maps derived from tractography round five but tested these probabilities in different ways. First, we performed a series of ANCOVAs that tested for influence on the volume percent (the fraction of suprathreshold voxels that favored striosome-like or matrix-like voxels, described in section 2.10). Collinearity among our insular subregion measures precluded our use of MANCOVA. The explanatory factors we considered for each subregion included compartment, hemisphere, handedness, sex, self-identified race, and age. Given the recent findings by Cabeen et al.,^46^ that hemispheric lateralization in the rostral-most insula is significantly influenced by sex, we also included the combined interaction of sex, hemisphere, and compartment. As we performed 19 separate ANCOVAs (one for each insular subregion), we set our significance threshold at p = 2.6×10^−3^ (0.05 / 19). Next, we assessed bias in structural connectivity at individual voxels across the whole insula with *randomise*, using the same parameters described for tractography round four, but masking the analysis using the whole-insula region we utilized as the seed for tractography. As this comparison was paralleled by our previously described iteration of *randomise*, we set a significance threshold of p = 0.025.

Finally, we performed a series of post-hoc comparisons to assess the influence of nucleus of origin or local striatal environment on compartment-specific biases (tractography rounds six and seven). For round six (testing the influence of nucleus of origin), we performed the same set of 19 ANCOVA tests that we utilized for tractography round five, so we set our significance threshold to equal our *a priori* ANCOVA comparisons (p = 2.6×10^−3^). We labelled all ANCOVA results that were <10 times the significance threshold (p = 2.6×10^−2^) as ‘trending’ towards significance. We assessed insulo-striate compartment selectivity, when projecting to location-shifted striatal masks (tractography round seven), using t-tests with a significance threshold of p = 2.6×10^−3^ (0.05 / 19).

## Results

### 3.1: Study Summary

We investigated structural connectivity between the insular cortex and the striatum to determine whether particular insular subregions were biased in their connectivity with the striatal compartments, striosome and matrix. First, we assessed the degree to which our striosome-like and matrix-like voxels matched the anatomic properties of striosome and matrix observed in animal and human tissue. Next, we mapped the insular subregions for compartment-level bias in structural connectivity. The first step in these analyses was to assess the location and amplitude of insulo-striate streamline bundles after they exited the insula. Next, we evaluated each insular subregion for compartment-specific bias in structural connectivity. At the levels of the whole insula, the insular subregions, and the subcortical white matter, insulo-striate structural connectivity with striosome-like voxels had markedly different location and amplitude than connectivity with matrix-like voxels. For some subregions, compartment bias was significantly influenced by a sex-by-hemisphere interaction. Finally, we tested alternate anatomic factors, unrelated to the striatal compartments, that might also influence insulo-striate bias. Neither nucleus-of-origin (caudate or putamen) nor local striatal environment was a significant or plausible explanatory factor for the compartment-specific biases we identified.

### 3.2: The relative location of striosome-like and matrix-like voxels

The typical location of striosome and matrix within the mammalian striatum is remarkably similar across species. In human^27,47,48^ and animal tissue,^24,26,27,49,50^ each compartment is arrayed in medio-lateral, rostro-caudal and dorsal-ventral gradients. Thus, while the striosomal labyrinth is present throughout the striatum, its branches are enriched in medial, rostral, and ventral sites. ^51^ Our connectivity-based parcellation method identified these same gradients: on average, striosome-like voxels are located more medial, more rostral, and more ventral than matrix-like voxels (Figure 1). As in striatal histology, one may find striosome-like or matrix-like voxels at any selected point in the striatum – but each compartment is biased in its most frequent locations. Note that these assessments of voxel location dealt with the voxels in our equal-volume masks (the most-biased voxels, filling the volume 1.5 standard deviations above the mean), not with every striosome-like and matrix-like voxel. Two factor ANOVA demonstrated that striatal compartment, the striatal nucleus of origin (caudate or putamen), and the interaction of these two factors all significantly influenced voxel location. In all planes, our compartment-like voxels matched the location biases demonstrated through prior histology assessments.

**Figure 1:**
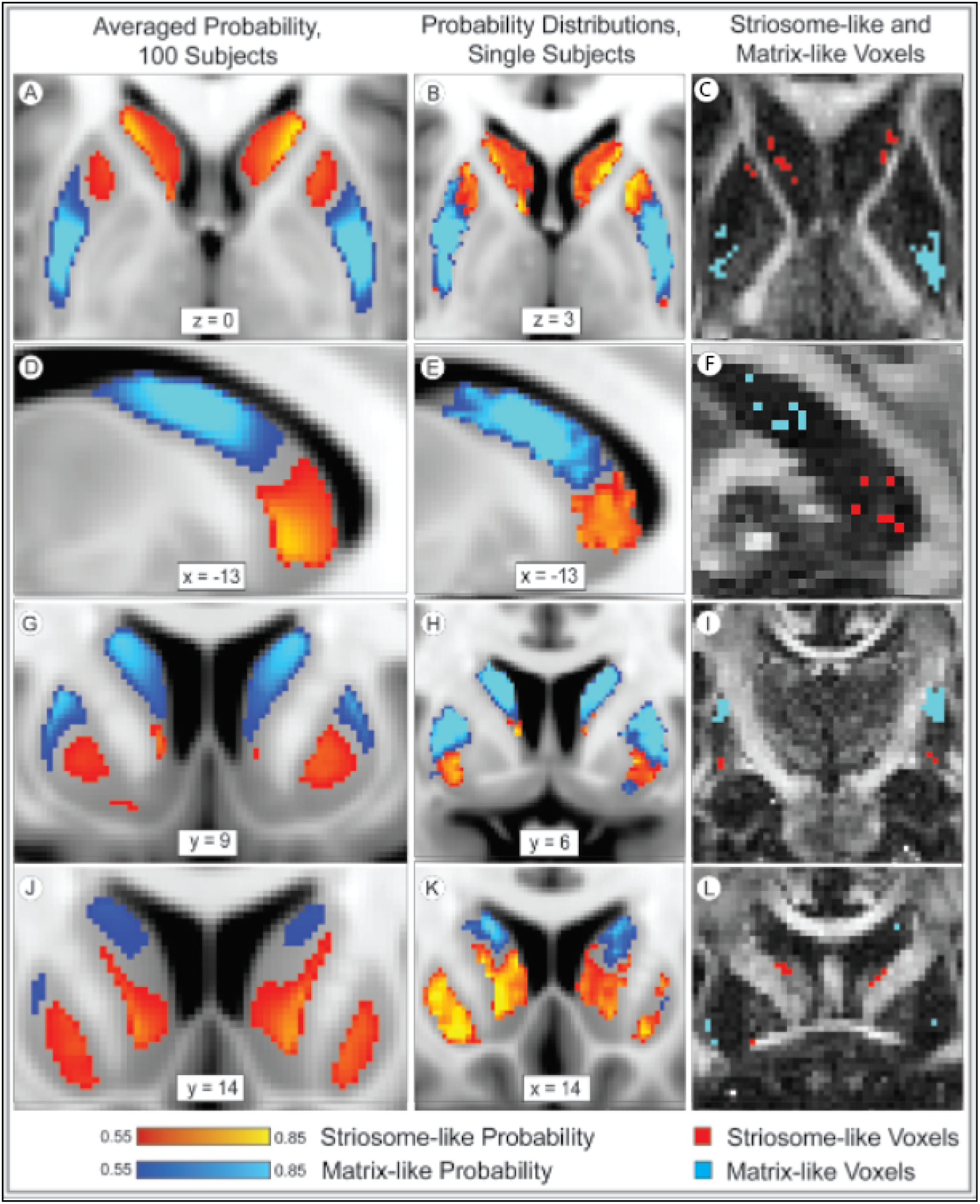
The distribution of striosome-like and matrix-like voxels matches the pattern expected from histology. The averaged probability distributions (left column) for striosome-like and matrix-like structural connectivity demonstrate that striosome-like probability (red-yellow) is enriched in the rostral, medial, and ventral striatum, while matrix-like probability (blue-light blue) is enriched in caudal, lateral, and dorsal striatum. This spatial pattern holds true for individuals as well (middle and right columns, all panels from the same subject). Striosome-like and matrix-like voxels (right column) are the most-biased voxels from each individual’s striosome-like and matrix-like probability distributions (middle column). The middle and right columns illustrate the same planes of section (in standard and native space, respectively) for this individual subject. Note that low-probability voxels (those near the 0.55 probability cut-off) are likely sampling a mix of striosome and matrix tissue. The highly biased voxels of the 1.5SD masks (right column) are substantially more specific and follow the dispersed pattern expected for striosome-like voxels. Probabilities are illustrated relative to the MNI_152_1mm standard in the left and middle columns. Compartment-like voxels (right column) are illustrated in native space (1.5 mm isotropic resolution) relative to that individual’s fractional anisotropy image. Coordinates follow MNI convention. Images follow radiographic convention.

The mean location of striosome-like voxels was 1.0 mm more medial (F_(1, 102)_=1,166; p = 1.2×10^−57^), 4.0 mm more rostral (F_(1, 102)_=1,629; p=1.6×10^−64^), and 6.2 mm more ventral (F_(1, 102)_=26,512; p = 4.5×10^−125^) than the mean location of matrix-like voxels. Origin within the caudate or putamen also had a significant influence on voxel location. Matrix-like voxels in the putamen were more lateral (F_(1, 102)_=29.3; p = 4.1×10^−7^), more caudal (F_(1, 102)_=139; p = 9.3×10^−21^), and more dorsal (F_(1, 102)_=20.7; p = 1.5×10^−5^) than matrix-like voxels in the caudate. Note that the relative positioning of each nucleus within the hemisphere did not drive these differences in location, as individual voxels were assessed relative to the centroid of their nucleus of origin. However, differences in the size and shape of the caudate and putamen may have allowed for more eccentric voxel placement in the putamen.

The interaction of compartment and nucleus of origin had a significant effect on voxel location in the y- (F_(1, 102)_=45.9; p = 8.2×10^−10^) and z-planes (F_(1, 102)_=285; p = 2.7×10^−31^). Simple main effects analysis of the compartment-nucleus interaction showed that in the y-plane, nucleus of origin had a significant effect on matrix location (F_(1, 66,709)_=146; p = 1.4×10^−33^), while in the z-plane, nucleus of origin had a significant effect on striosome location (F_(1, 66,709)_=286, p = 5.0×10^−63^).

### 3.3: The relative abundance of striosome-like and matrix-like voxels

In human tissue, striatal MSNs are present in a ratio of approximately 15% striosome, 85% matrix.^48^ This ratio is relatively constant across all mammalian species reported to date, despite the many-fold increase in both brain size and total striatal volume from rodent to human. Of voxels that were highly biased (P ≥ 0.87), striosome-like voxels comprised 12.2% and matrix-like voxels comprised 87.8% of striatal volume (SEM, ±0.68%). The volume of the striosome-like and matrix-like compartments was not meaningfully different from the ratio expected from histology.

### 3.4: Compartment-specific biases in corticostriate structural connectivity

Injected tract tracers in animals demonstrated that axons originating in primary motor cortex project almost exclusively to the matrix.^24,52–54^ Similarly, axons from posterior orbitofrontal cortex are strongly biased toward the striosome.^25,40,55^ We previously demonstrated that parcellating the human striatum based on differential connectivity replicates these and other patterns of bias originally established in animals, though in different subjects and using a different diffusion imaging protocol.^30^ We performed N-1 (“leave one out,” tractography round two) striatal parcellations, generated 1.5SD masks from these parcellations, and then assessed compartment-selective connectivity bias using the left-out regions as seeds and compartment-like voxels as targets (tractography round three). We quantified all voxels within the cortical seed region with high probability of connection (P≥0.87) to either compartment-like mask. In the present cohort, 95.4% of biased primary motor cortex voxels favored matrix-like striatal voxels (p = 4.2×10^−85^). Similarly, 98.3% of biased voxels in the posterior orbitofrontal cortex favored striosome-like voxels (p = 1.1×10^−90^).

Parcellating striatal voxels based on differential structural connectivity replicates many anatomic features of the striosome and matrix: our striosome-like and matrix-like voxels match the spatial distribution, relative abundance, and region-specific biases in connectivity expected from striatal histology. However, we urge readers to bear in mind that we have identified voxels that share many properties of the striatal compartments, but these indirect and probabilistic parcellations are not the equivalent of directly identifying striosome and matrix tissue.

### 3.5: Mapping insulo-striate streamlines

We compared the amplitude and location of streamline bundles that linked the insula and either striosome-like or matrix-like striatal voxels (tractography round four; Figure 2). Total streamlines contacting matrix-like voxels did not differ between left and right hemispheres (right, +10.6%, p = 0.11). Streamlines that contacted striosome-like voxels were divided into rostral and caudal bundles, which we assessed separately. Total streamlines in the rostral striosome-favoring bundle were substantially more abundant in the left hemisphere (2.6-fold larger; p = 5.9×10^−11^). In the caudal striosome-favoring bundle total streamlines did not significantly differ between the hemispheres (right, +33.2%; p = 0.31). Note that the mean insular mask volume differed between the hemispheres by only 48 voxels (1.9% of total insular volume). This small insular asymmetry is not a plausible cause of the large, compartment- and location-specific differences between the hemispheres. Streamlines that contacted striosome-like voxels were 2.7-fold more abundant than those that contacted matrix-like voxels (striosome-like: 1.1×10^5^, matrix-like: 4.1×10^4^; p = 1.2×10^−11^).

**Figure 2:**
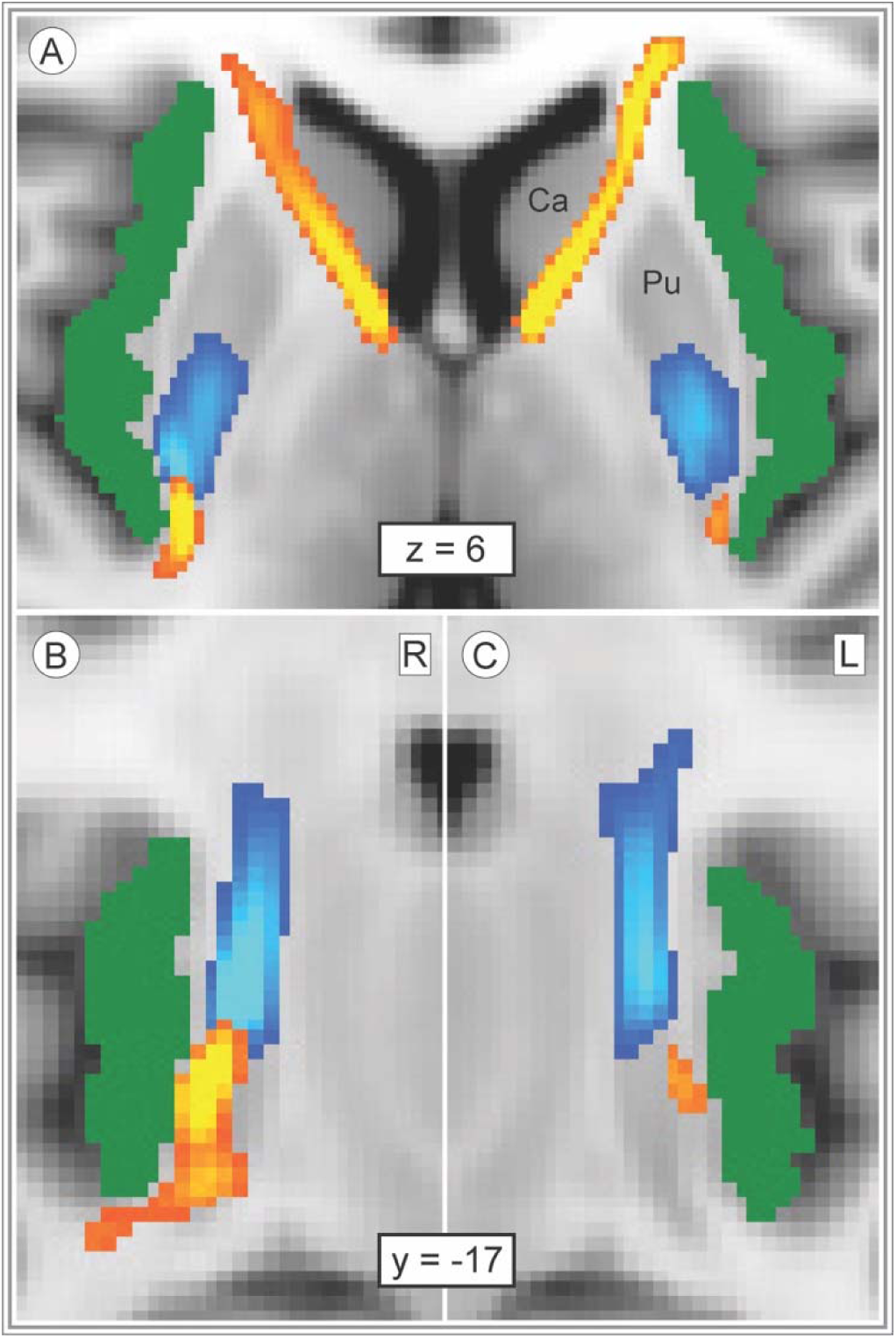
Insulo-striate streamlines followed distinct paths to reach striosome-like vs. matrix-like voxels. Streamlines bound for striosome-like (red-yellow) or matrix-like (blue-light blue) voxels are illustrated in the axial (A) or coronal plane (B, right; C, left; normalized amplitude: 0.5-1.0. Our whole-insula mask is shown in green. Striosome-bound and matrix-bound streamlines followed highly similar paths in the left and right hemispheres, though we generated tractography in each hemisphere independently. Streamlines that originated in the rostral insula that targeted striosome-like voxels exited the insula laterally and traversed the corona radiata and anterior limb of the internal capsule (A). Streamlines that targeted matrix-like voxels, and caudoventral striosome-bound streamlines, exited the insula medially and predominantly transited via the external capsule (A-C). Coordinates follow MNI convention. Images follow radiographic convention. Abbreviation: Ca, Caudate; L, left; Pu, Putamen; R, right.

Next, we assessed the locations of compartment-specific insulo-striate streamlines within the subcortical white matter (Figure 2). Though we performed tractography independently in each hemisphere, the location of streamline bundles was highly similar in the left and right hemispheres (Fig. 2 A-C). Streamlines bound for matrix-like voxels exited the insula in a single bundle and appeared to transit within the external or extreme capsules (difficult to distinguish at this resolution). In contrast, streamlines bound for striosome-like voxels transited within distinct rostral and caudal bundles (Fig. 2 C-E). While the caudal striosome-bound bundle exited the insula medially and transited the extreme/external capsule (paralleling the matrix-bound bundle, Fig. 2 A-C), the rostral striosome-bound bundle exited the insula from the rostro-lateral surface and transited within the corona radiata and anterior limb of the internal capsule. The rostral striosome-bound bundle appeared to primarily target the caudate, while the posterior striosome-bound bundle appeared to primarily target the putamen.

Insulo-striate streamlines that contacted striosome-like voxels were spatially segregated from streamlines that contacted matrix-like voxels. We isolated the core of each streamline bundle (retaining the uppermost 25%, 50%, or 75% of the streamline bundles, thresholds we investigated previously; ^44^ and assessed the overlap of the core striosome-bound and matrix-bound bundles. The uppermost 25% and 50% of striosome-bound and matrix-bound bundles had no voxels overlapping in either hemisphere. Bundles that included the uppermost 75% of voxels had no overlap in the left hemisphere, and only 17 voxels overlapped in the right hemisphere (DSC = 0.0004).

Striosome-favoring and matrix-favoring streamlines were highly segregated when assessed at a voxelwise level, without amplitude thresholds, as well (FSL *randomise*). Following stringent familywise-error and Bonferroni correction, 66.8% of the voxels in our subcortical bounding mask were significantly biased towards one of the two compartments. These significant clusters closely approximated the mean distributions seen in Fig. 2 A-C. The striosome-favoring distribution occupied a larger fraction of subcortical voxels (47.1%) than the matrix-favoring distribution (19.7%). The streamline density within these significant clusters underscored the marked difference in subcortical location for compartment-specific streamlines: within the striosome-favoring significant clusters (60,432 voxels), 9,161 streamlines/voxel contacted striosome-like voxels, while 305 streamlines/voxel contacted matrix-like voxels; within the matrix-favoring significant cluster (25,319 voxels), 318 streamlines/voxel contacted striosome-like voxels, while 3,831 streamlines/voxel contacted matrix-like voxels. 3.5: Spatial distribution of compartment-specific connectivity biases within the insula

To map connectivity for each insular subregion, we assessed insulo-striate connectivity at each insular voxel using classification targets mode (tractography round five). Insular voxels that seeded streamlines bound for striosome-like voxels were largely segregated from insular voxels whose streamlines reached matrix-like voxels (Figure 3). We calculated the root-mean-square (RMS) distance between the centers of gravity (COG) for striosome-favoring and matrix-favoring probability distributions for each individual and hemisphere. Given the visible separation of rostral and caudal striosome-favoring probability clusters (Fig. 3), we chose to compare matrix COG with the rostral striosome COG, and matrix COG with the caudal striosome COG. The matrix-favoring cluster was significantly closer to the centroid of the insula than both striosome-favoring clusters (rostral striosome-favoring cluster: 11.0 mm greater RMS distance, p = 1.9×10^−118^; caudal striosome-favoring cluster: 8.5 mm greater RMS distance, p = 6.1×10^−111^). Relative to the matrix-favoring cluster, the rostral striosome-favoring probability cluster was 16.8 mm more rostral, 2.0 mm more medial, and 7.7 mm more ventral within the insula (p-values, respectively: 1.1×10^−109^, 1.3×10^−36^, 2.2×10^−67^). Relative to the matrix-favoring cluster, the caudal striosome-favoring probability cluster was 11.7 mm more caudal, 2.4 mm more lateral, and 7.0 mm more ventral within the insula (p-values, respectively: 2.4×10^−84^, 6.4×10^−82^, 2.1×10^−60^).

**Figure 3:**
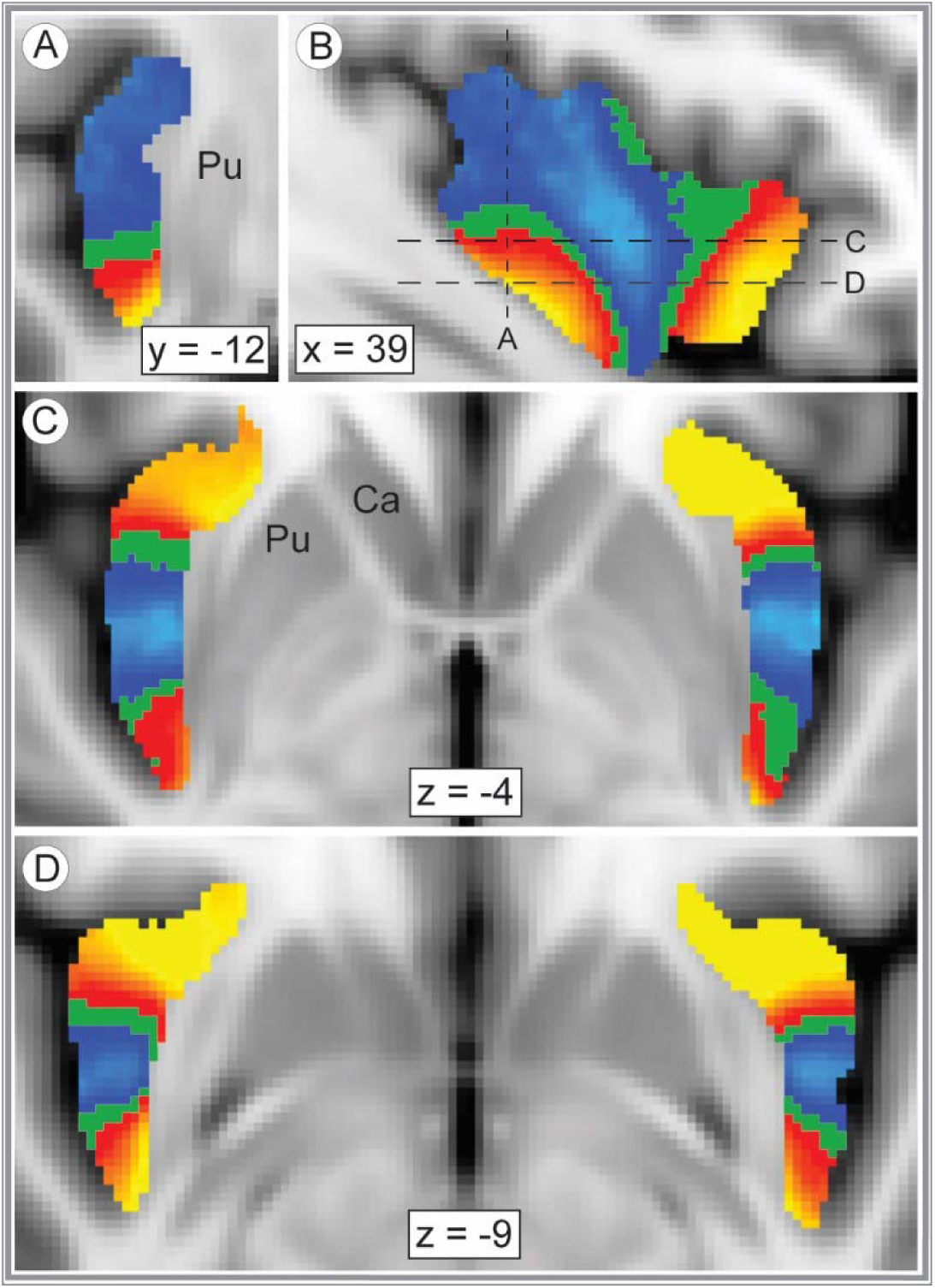
Insulo-striate connectivity with the striosome- or matrix-like compartments is highly segregated. Insular voxels whose connectivity favored striosome-like voxels (red-yellow) or matrix-like voxels (blue-light blue) are overlaid on the insular seed mask (green). All voxels were visualized with probability thresholds of 0.55–0.85, revealing the insular mask in minimally-biased (P < 0.55) voxels. The highest connection probabilities are shown in yellow and light blue for striosome and matrix-like connectivity, respectively. Coronal (A, right hemisphere), sagittal (B, right hemisphere), and axial (C, D) views reveal the distinct zones of insulo-striate structural connectivity. The planes of visualization in A, C, and D are represented by dashed lines in B. Coordinates follow MNI convention. Images follow radiographic convention. Abbreviation: Ca, Caudate; Pu, Putamen.

### 3.6: Compartment-specific biases among the insular subregions

Next, we quantified compartment-specific connectivity (tractography round five) for each insular subregion in each hemisphere (Figure 4, Supplemental Video 1). For every subregion, compartment-specific connectivity biases matched in the left and right hemispheres. Therefore, we included both hemispheres in subsequent assessments of connectivity bias. We quantified compartment-specific connectivity in two ways. First, we extracted volume (the number of voxels with probability ≥ 0.55 for striosome- or matrix-favoring connectivity) within each subregion and assessed the influence of striatal compartment on connectivity using ANCOVA, with a range of demographic and anatomic factors as covariates (detailed in Section 2.11). This approach investigated the influences on compartment-specific structural connectivity <u>within biased voxels.</u> Second, we performed voxelwise, non-parametric comparisons (striosome-favoring vs. matrix favoring probability distributions, using *randomise*) to identify voxels with significant biases in structural connectivity (familywise error corrected for multiple comparisons). Within each insular subregion we then extracted the number of significantly biased voxels <u>as a fraction of all voxels in that subregion</u>. Comparing the results of these two approaches provided anatomic detail not evident in single measures of mean connectivity across a subregion.

**Figure 4:**
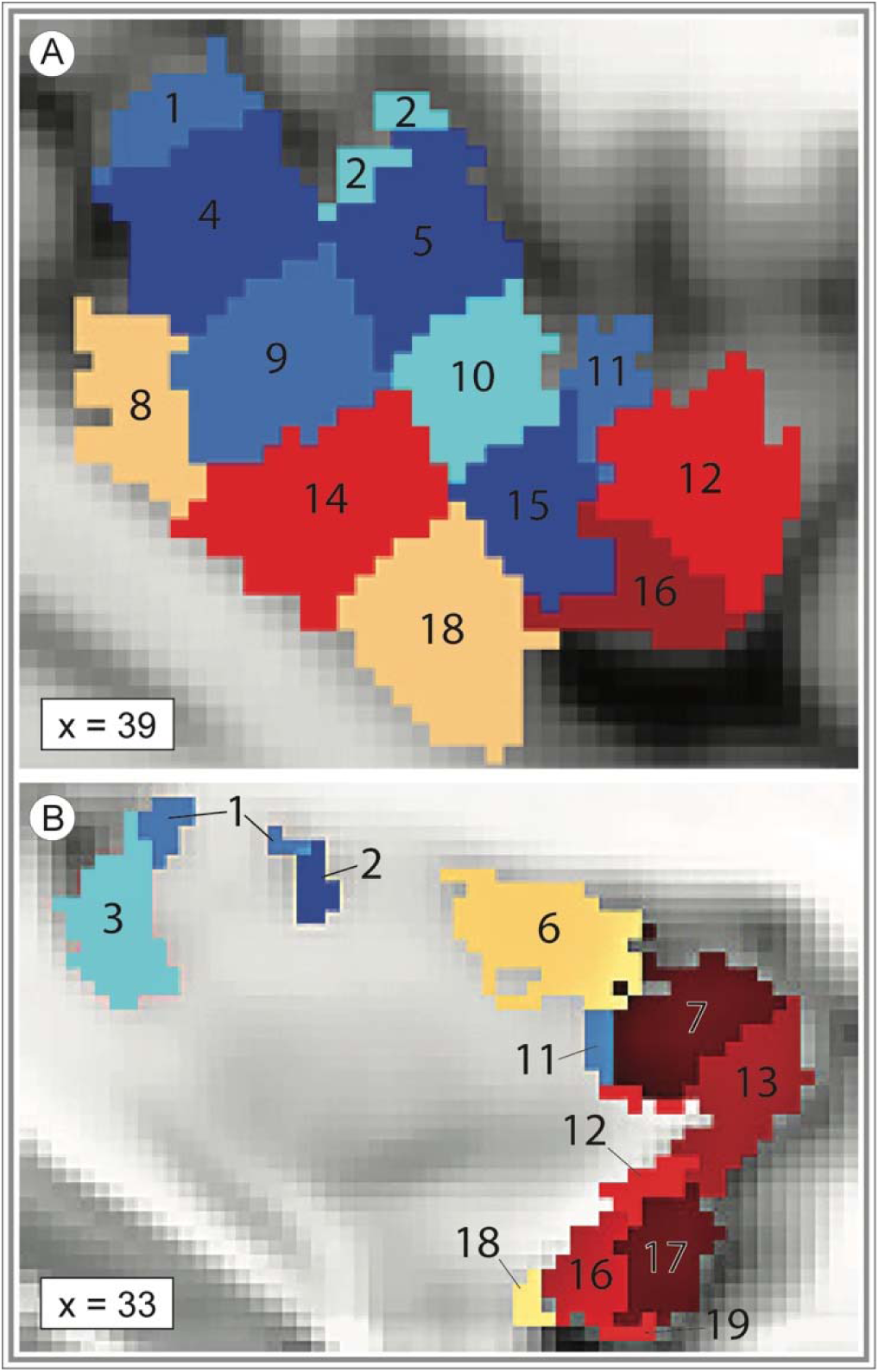
Bias in striatal connectivity clusters among neighboring insular subregions. The 19 masks utilized to extract subregion-specific connectivity estimates are superimposed on the MNI152_T1_1mm template brain. The ROIs shaded red represent striosome-biased insular subregions and are generally more rostral and ventral. The ROIs shaded blue represent matrix-biased subregions and are generally more caudal and dorsal. The variation in color of the ROIs within each red or blue region is only to distinguish ROI subregions – shading does not indicate the magnitude of bias towards either compartment. The ROIs shaded tan represent subregions with no compartment-specific bias. ROI numbers correspond to those in the bar graph, Figure 5. The sagittal planes of visualization (x = 39 or 33) are lateral (A) and medial (B) views, respectively, of the left insula. Coordinates follow MNI convention.

The majority of insular subregions (16 of 19, 84%) were significantly biased towards either striosome-like or matrix-like voxels (Figure 5, Table 2). In every case, the biases identified through ANCOVA and voxelwise testing concurred. Nine subregions were significantly biased towards matrix-like compartments: subregions 1, 2, 3, 4, 5, 9, 10, 11, 15. Seven subregions were significantly biased towards striosome-like voxels: subregions 7, 12, 13, 14, 16, 17, 19. Three subregions had no significant bias in structural connectivity: subregions 6, 8 and 18. Insular subregions with bias towards striosome-like voxels tended to be in rostral and ventral subregions, while those with bias towards matrix-like voxels tended to be in caudal and dorsal subregions (Figure 4). Subregions without significant bias straddled the boundaries between striosome- and matrix-favoring insular zones, and thus included large numbers of voxels that were biased toward either compartment (Figure 5, purple violin plots).

**Figure 5:**
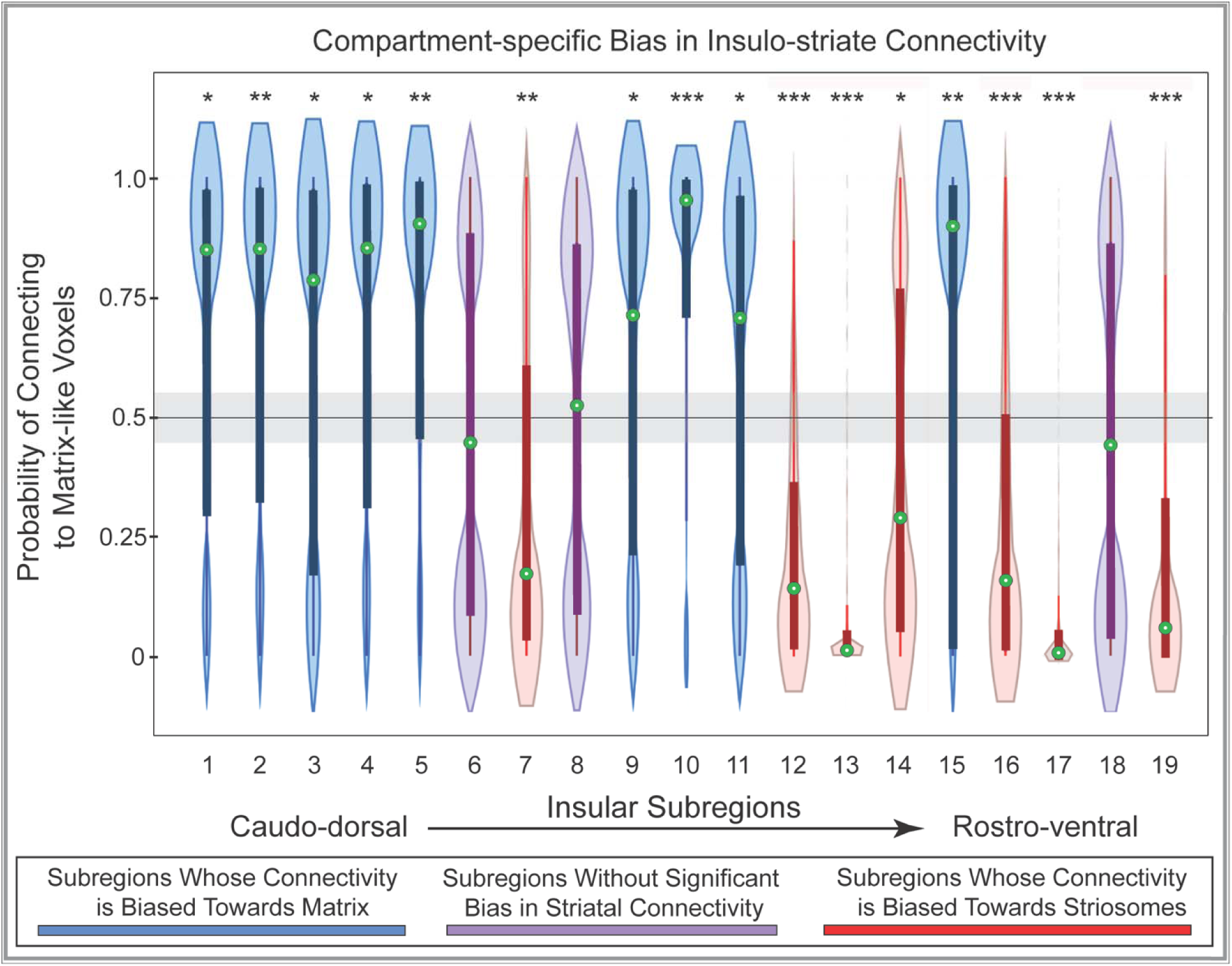
Structural connectivity between 19 insular subregions and the striatal compartments is largely biased toward either striosome-like or matrix-like voxels. Insular subregions are presented from the caudo-dorsal to the rostro-ventral insula (left-to-right). Values higher than 0.55 on the y-axis (above the grey bar) indicate matrix-like connectivity bias; values lower than 0.45 (below the grey bar) indicate striosome-like connectivity bias. Nine insular subregions were significantly biased towards matrix-like voxels (blue violin plots), seven subregions were significantly biased towards striosome-like voxels (red violin plots), and three subregions had no significant compartment bias (purple violin plots). Green circles indicate the median for that subregion. Box and whisker plots are superimposed on each violin plot. The values for matrix-like and striosome-like connectivity for any individual insular subregion always sum to one. Therefore, we present values only for matrix-like connectivity for each insular subregion. Significance thresholds: *, p = 2.6×10^−3^; **, p = 2.6×10^−9^; ***, p = 2.6×10^−15^.

**Table 2:** Compartment-Specific Biases in the Insular Subregions. Each insular subregion is numbered to match the insular subregion masks defined by Ghaziri et al. (2018; illustrated in Figure 4). “Compartment Bias” indicates the compartment that is the primary target of streamlines seeded from within that subregion. ANCOVA testing (middle three columns) assessed compartment-biased voxels for factors that influenced that bias. For these tests, percentages from 55 to 100% indicate a progressively increasing bias toward matrix-like voxels, and percentages from 45 to 0% indicate a bias toward striosome-like voxels. Voxelwise testing (*randomise*, right two columns) assessed all insular voxels for significant bias (familywise error-corrected significance threshold, p = 0.025), expressed as the percent of a subregion’s total volume that was significantly biased toward either compartment. While some subregions included significant voxels for both compartments (e.g., subregion 14), 10 of 19 subregions included only voxels biased toward one compartment. Note that subregions 6, 8, and 18 were not significantly biased and thus are not included here.

| Insular Subregion | Compartment Bias | Results of ANCOVA testing |  |  | Results of *randomise* testing |  |
| --- | --- | --- | --- | --- | --- | --- |
| | | % of biased voxels favoring matrix | F statistic ($F_{1,379}$) | p-value | % of subregion voxels with matrix bias | % of subregion voxels with striosome bias |
| Subregion 1 | Matrix-like | 65.9 | 71.2 | $p = 6.9 \times 10^{-16}$ | 63.6 | 0.0 |
| Subregion 2 | Matrix-like | 66.9 | 71.2 | $p = 4.1 \times 10^{-18}$ | 85.4 | 0.0 |
| Subregion 3 | Matrix-like | 61.4 | 33.0 | $p = 1.9 \times 10^{-8}$ | 32.5 | 0.0 |
| Subregion 4 | Matrix-like | 65.0 | 61.7 | $p = 4.3 \times 10^{-14}$ | 70.1 | 0.0 |
| Subregion 5 | Matrix-like | 71.0 | 144.3 | $p = 2.2 \times 10^{-28}$ | 82.7 | 0.0 |
| Subregion 7 | Striosome-like | 33.6 | 91.4 | $p = 1.5 \times 10^{-19}$ | 0.0 | 63.4 |
| Subregion 9 | Matrix-like | 60.0 | 26.8 | $p = 3.7 \times 10^{-7}$ | 50.7 | 2.9 |
| Subregion 10 | Matrix-like | 76.7 | 231 | $p = 5.0 \times 10^{-41}$ | 93.6 | 0.0 |
| Subregion 11 | Matrix-like | 58.8 | 22.6 | $p = 2.8 \times 10^{-6}$ | 51.0 | 0.18 |
| Subregion 12 | Striosome-like | 21.9 | 542 | $p = 4.9 \times 10^{-75}$ | 1.3 | 74.6 |
| Subregion 13 | Striosome-like | 9.3 | 1,723 | $p = 4.9 \times 10^{-143}$ | 0.0 | 98.4 |
| Subregion 14 | Striosome-like | 40.4 | 27.8 | $p = 2.2 \times 10^{-7}$ | 16.3 | 43.3 |
| Subregion 15 | Matrix-like | 68.4 | 88.9 | $p = 4.2 \times 10^{-19}$ | 79.4 | 1.9 |
| Subregion 16 | Striosome-like | 28.9 | 178 | $p = 1.7 \times 10^{-33}$ | 5.6 | 78.0 |
| Subregion 17 | Striosome-like | 7.3 | 2,797 | $p = 5.0 \times 10^{-177}$ | 0.0 | 97.2 |
| Subregion 19 | Striosome-like | 21.1 | 403 | $p = 1.4 \times 10^{-61}$ | 0.0 | 82.8 |

Supplemental Video 1: A video illustrating the left insula, our whole-insula seed mask, and the insular subregion masks that we used to map compartment-specific bias in insulo-striate connectivity. Subregion masks were generated by Ghaziri et al., 2018.

Note that comparing the results of ANCOVA and voxelwise testing is useful to distinguish between punctate areas of bias and broader patterns of bias that involve most or all of a subregion. For example, ANCOVA testing (the percent of biased voxels) yielded similar results for subregions 2 and 3, with roughly 2:1 biased voxels (matrix:striosome). Assessing the results of voxelwise testing, however, demonstrates that 85.4% of all voxels in subregion 2 were biased toward matrix-like targets, while in subregion 3 only 32.5% of voxelswere significantly biased toward matrix-like targets – and 67.5% of subregion 3 voxels were not significantly biased toward either compartment. Therefore, while streamlines seeded in subregions 2 and 3 were both significantly biased toward matrix-like voxels, this pattern is much more broadly-based in subregion 2.

We included demographic and experimental variables in our ANCOVA testing to determine the impact of these factors on compartment-specific bias in insulo-striate connectivity. When tested in isolation, age, sex, self-identified race, handedness, and the hemisphere of origin did not have a significant influence on compartment-specific bias, for any region. However, given the recent findings by Cabeen et al.^46^ that the histologic structure of the frontal insula is partially determined by hemisphere and subject sex – we also tested for the interaction of sex and hemisphere on compartment-specific bias. Of the seven rostral subregions that approximated the frontal insula (as mapped by Cabeen et al.) the sex-by-hemisphere interaction was a significant contributor to compartment bias in three subregions: subregion 11, *F_(4,379)_*= 8.2, p = 2.2×10^−6^; subregion 13, *F_(4,379)_* = 15.4, p <1.2×10^−11^; subregion 19, *F_(4,379)_*= 8.8, p = 8.0×10^−7^. For three additional rostral subregions, the sex-by-hemisphere interaction trended toward significance: subregion 7, *F_(4,379)_* = 3.7, p = 5.3×10^−3^; subregion 12, *F_(4,379)_*= 3.1, p = 1.6×10^−2^; subregion 17, *F_(4,379)_*= 3.0, p = 1.7×10^−2^. In contrast, for the 12 regions that did not approximate the frontal insula, no region reached or trended towards significance.

### 3.7: The influence of nucleus of origin and rostro-caudal position on insulo-striate connectivity

While compartment-specific biases may explain the spatially segregated patterns of connectivity we described, alternate anatomic factors may influence insulo-striate connectivity as well. We considered the hypothesis that the nucleus of origin – whether a striosome-like or matrix-like voxel resided in the caudate or putamen – influenced the strength or specificity of compartment-specific insular subregion bias. Using the output of tractography round one (striatal parcellation), we generated a second round of striosome-like and matrix-like masks that were proportional to the relative volume of the caudate and putamen. That is, we selected voxels based on the strength of the bias in connectivity, but rather than selecting the 180 voxels with the largest compartment-specific biases (as we utilized in tractography rounds one through five), caudate and putamen each had a quota of highly biased voxels based on the relative volume of caudate and putamen. Therefore, these striatal masks were biased in their connectivity, but not as biased as those whose sole criteria for selection was connectivity, independent of whether they resided in caudate or putamen. We then completed insulo-striate CTT with these proportionate striatal masks as the targets for CTT (tractography round six). Relative to tractography round five, no subregion began with bias toward one compartment and shifted to bias towards the opposite compartment. When ranked by the amplitude of bias, biased subregions in the proportionate masks condition followed the same order as observed in the original 1.5SD masks – with one exception, in which two subregions differed by 1% in the original and proportionate masks conditions. With proportionate striatal masks, all insular subregions were slightly more biased towards striosome-like voxels (shifting by 0.92–9.2%, mean: 5.8% shift). This change to proportionate target masks led subregions 3 and 9 to fall from slight matrix-bias (Figure 5) to neutral. No subregion shifted to greater matrix-biased connectivity. One insular subregion was significantly more biased toward striosome-like voxels with proportionate masks (subregion 19, shifting from 21.1% matrix bias to 11.9%, p = 6.2×10^−4^) – all other shifts in bias toward striosome-like voxels were non-significant. We conclude that the magnitude of compartment-specific bias was a more important driver of insulo-striate connectivity than whether a compartment-like voxel was found in caudate or putamen.

The insulo-striate biases we identified share a spatial organization with the striatum: both striosome-like voxels and striosome-favoring insulo-striate projection are enriched in the rostral and ventral portions of their region, with matrix-favoring and matrix-like voxels enriched centrally. Therefore, we considered the possibility that these biases reflect simple spatial ordering of streamlines, an effect of the “neighborhood” in which a voxel resides rather than its compartment-specific bias. We addressed this possibility by jittering the voxel location of the voxels of our striosome-like and matrix-like masks and then compared our initial tractography (which targeted our precisely-selected compartment-like masks) with tractography that targeted these location-shifted voxels. Notably, we shifted the location of each voxel individually, and by a random distance (<u>±</u>0-3 voxels) in each plane. The mean location shift for individual voxels was 1.9 voxels (range: 0-3 voxels) in each plane. However, as the direction of shift was random for each voxel and in each plane, the mean location shift was negligible (<1 mm difference for each plane - x: 0.16 mm shift, y: 0.10 mm shift, z: 0.79 mm shift). The RMS distance from the centroid of either caudate or putamen was 7.6 mm for our starting compartment-like masks (6.6 mm for matrix, 8.7 mm for striosome), and 7.4 mm for the randomly shifted voxels (7.3 mm for matrix, 7.5 mm for striosome). The proximity of these location-shifted voxels allowed us to distinguish compartment-specific and “neighborhood” influences on insulo-striate connectivity.

Our striosome-like and matrix-like masks had mean connectivity biases of 0.84 and 0.97, respectively (neutral: P<0.55; complete bias: P=1.0). In contrast, our location-shifted voxels had compartment biases of 0.56 for striosome-adjacent and 0.84 for matrix-adjacent masks. The fact that mean bias changed substantially more in striosome-adjacent than in matrix-adjacent voxels matches expectations based on striatal tissue - since 85% of striatal volume lies within the matrix (Johnston et al. 1990), shifting location at random is substantially more likely to locate a voxel occupied by matrix. Shifting the location of a highly-biased matrix-like voxel will likely select a less-biased, but still matrix-like, voxel. In contrast, shifting the location of striosome-like voxels may select a less-biased striosome-like voxel (since their location is relatively clustered) or an abundant matrix-like voxel. Our striosome-adjacent voxels reflected this near-neutral pattern of bias. When tractography targeted these location-shifted voxels (tractography round seven), connectivity was markedly and significantly shifted toward matrix-adjacent masks for all 19 insular subregions (range of increase in bias across 19 subregions: 0.60-0.98, 0.21-0.56 greater than prior matrix-like bias; for comparisons of precise-to-shifted targets, p-values ranged from 4.7×10^−16^ – 2.7×10^−56^). Though individual compartment-adjacent voxels were shifted <2 voxels from their precisely-selected starting points, and the mean location of these location shifts was <1 mm, the striosome-favoring insular subregions lost all bias toward striosome-adjacent voxels. The biases we identified in insulo-striate projections were driven by the compartment-specific differences in structural connectivity, not by their relative “neighborhood” within the striatum.

## Discussion

### 4.1: Overview of Results

We assessed structural connectivity in 100 healthy human subjects to determine whether and where insulo-striate projections are biased toward one of the two striatal compartments, striosome or matrix. Voxels with striosome-like or matrix-like structural connectivity matched the anatomic properties of striosome and matrix demonstrated through histology: their typical locations within the striatum, relative abundance, somatotopy, and the compartment-specificity of cortico-striate projections. Insulo-striate streamline bundles followed segregated paths to reach the striatum, depending on if they targeted striosome-like or matrix-like voxels. We found that insular subregions whose streamlines primarily contacted striosome-like voxels were spatially segregated from subregions whose streamlines primarily contacted matrix-like voxels: 16 of 19 insular subregions had a significant bias toward one striatal compartment. Though we generated tractography independently in the left and right hemispheres, these subregion biases and the locations of compartment-specific streamline bundles were highly similar between hemispheres. Biases in insulo-striate structural connectivity were specific for the precisely selected striatal voxels in our striosome-like and matrix-like masks and these biases were not shared by voxels in their immediate vicinity. We compared our quantitative and qualitative structural connectivity results with prior histologic and imaging studies in human, primate, and non-primate animal studies. Our diffusion MRI-based results matched the compartment-specific biases in structure and function described in multiple species through diverse experimental methods.

### 4.2: Limitations of Connectivity-based Striatal Parcellation

While probabilistic tractography is a powerful tool for understanding structural connectivity non-invasively in living organisms, this technique has notable limitations. Tractography cannot distinguish between afferent and efferent connections, cannot detect synapses, and is limited by the millimeter-scale resolution of diffusion MRI signal acquisition. Image resolution is a specific and important limitation of our parcellation method: the size mismatch between striosome branches (0.5-1.25 mm in diameter in coronal sections)^27,48^ and our diffusion voxels (1.5 mm isotropic) assures that each striosome-like voxel includes some fraction of matrix tissue. Therefore, we restricted our quantitative assessments to only the most-biased voxels, those whose connection bias was ≥1.5 standard deviations above the mean. While this approach replicated many anatomic features of striosome and matrix – their spatial distribution, relative abundance, somatotopy, and their patterns of extra-striate connectivity – readers should bear in mind that our method did not directly identify striosome and matrix tissue. Higher resolution diffusion acquisitions will reduce, but not eliminate, this mixed-sampling limitation.

Imperfections in the segmentation or registration of our regions of interest and bounding masks (discussed in Methods) may have reduced the anatomical precision of our tractography. However, the test-retest accuracy of our histologically-based striatal parcellation method (99.8%)^30^ suggests that such imperfections have a small influence on parcellations. Another potential source of error in our approach is the coexistence of MSN populations and *en passant* white matter bundles within striatal voxels. Jones and Cercignani^58^ estimated that up to 90% of brain voxels may contain multiple fiber pathways; this is especially problematic in the striatum, with its intercalated descending cortical projections. For this reason, we chose to model crossing fibers using the FSL tool *bedpostX*, which estimates tensors for each of three principal diffusion directions. Notably, alternate models for estimating diffusion fibers may perform better than the tensor-based tools we utilized, including models that utilize fiber orientation distribution functions^59^ or Q-ball imaging,^60^ available through the widely-adopted freeware tools MRtrix and DSI Studio, respectively. While these limitations to probabilistic tractography must be considered, we found in every point of comparison that our tractography-based results were consistent with prior histologic and MRI-based assessments of the insula and the striatal compartments. Finally, the insular segmentations we utilized are not the only reasonable method for subdividing this structure. Researchers interested in alternative insular subregions could extract measures of connectivity bias from our existing insular probability maps, tailoring these methods to their own investigations.

There is substantial evidence that cortical regions and sub-portions of subcortical nuclei are involved in distinct functional and structural networks. For example, patients with lesions in the pulvinar nucleus of the thalamus exhibited chronic neuropathic pain, whereas patients with lesions concentrated in adjacent thalamic nuclei had no pain symptoms.^61^ Electrical stimulation of the dorso-caudal insula produced somatic pain, with insular somatotopy corresponding to specific body sites.^62^ Insular functional activation increases following heat-induced^2^ and capsaicin-induced pain^63^ specifically in this same dorso-caudal insular site. In both the anterior pulvinar^39^ and dorso-caudal insula, structural connectivity is significantly biased toward matrix-like voxels. Whether the dorso-caudal insula and anterior pulvinar nucleus act in concert to mediate pain in humans, with the matrix as an obligate network hub, will require future investigation. Insulo-thalamic connectivity has been shown to operate within distinct CSTC loops.^64^ The anterior insula operates in a discrete regulatory circuit with the dorsal striatum and mediodorsal thalamus across a broad range of psychiatric disorders.^65^ Similarly, reward cues activate the ventral striatum, rostral insula, and caudal thalamus (as a group), but non-reward cues do not.^66^ The rostral orbitofrontal cortex shares a distinct functional connectivity network with the insula and mediodorsal thalamus.^67^ The functional connectivity between these insular subregions and diverse brain areas supports the premise that insular subregions may operate within distinct CSTC loops. The highly segregated routes followed by striosome-bound and matrix-bound insulo-striate streamline bundles (Fig. 2) underscore the premise that insular subregions may be embedded within distinct CSTC loops, mediated in part through segregation by striatal compartment.

### 4.3: Network implications of segregated insulo-striate connectivity

Prior studies that injected anterograde tracers in the agranular and frontoinsular cortex of the cat, macaque, or mouse demonstrated preferential projections to the striosome, with minor projections to the matrix. ^25,29,68^ Our results in humans concur with these histologic studies: streamlines seeded in the rostral and ventral portions of the insula predominantly reach striosome-like voxels. Although direct compartment-specific connectivity of the dorsal and caudal portions of the insula has not been mapped using injected tracers in animals, to the best of our knowledge, cortical regions that were previously shown to be matrix-favoring also selectively projected to the caudal insula.^6–8^ The present results, based on bait regions that were validated through animal tract tracing studies and tested in a large, broadly representative neuroimaging cohort, confirm and extend these prior findings, and suggest that compartment-specific biases in insulo-striate connectivity cluster by insular subregions (Figures 2, 3).

### 4.4: Compartment-specific connectivity and insular cytoarchitecture are spatially similar

Compartment-specific projections may be related to underlying differences in the cytoarchitecture of the rostral vs. caudal, and ventral vs. dorsal insula (Figure 6). Cell density and granularity increases from the rostroventral to caudodorsal insula, generally in a wave-like pattern (Figure 5).^5,69,70^ This cytoarchitectural segregation of the insula aligns with the compartment-specific biases in structural connectivity we observed. The rostral agranular area, as well as the central dysgranular area that extends along the ventral rostro-caudal axis of the insula, are spatially colocalized with our rostral and caudo-ventral zones that preferentially project to striosome-like targets.^5,71^ The similarity in cytoarchitecture (agranularity) between the rostral insula and the ventral portion of the dorsal insula (Figure 6) overlaps the insular zones we identified as significantly biased towards striosome-like voxels (Figure 4). Intriguingly, a second insular gradient, the abundance of von Economo neurons (VENs), also parallels the biases in insulo-striate connectivity we observed (Figure 5).^72,73^ VENs are highly enriched in humans and hominid primates, relative to other mammals. VENs are restricted to a few cortical sites, including the rostral insula.^74,75^ Though the functions of VENs are incompletely understood, they are proposed to play an outsized role in social-emotional cognition. ^74–76^ Insular VENs are most abundant in rostroventral subregions and dwindle as one moves caudally and dorsally, similar to the striosome-favoring bias in structural connectivity seen here. No prior studies mapped VEN projections with the striatal compartments, to the best of our knowledge. However, a recent report found that *Lypd1*, a peptide marker of VENs,^77^ is also selectively enriched in the striosome.^78^ If future studies demonstrate that VENs selectively project to the striosome, this association may serve as a substrate for limbic symptoms in compartment-selective neurodegenerative diseases.^16,81^

**Figure 6:**
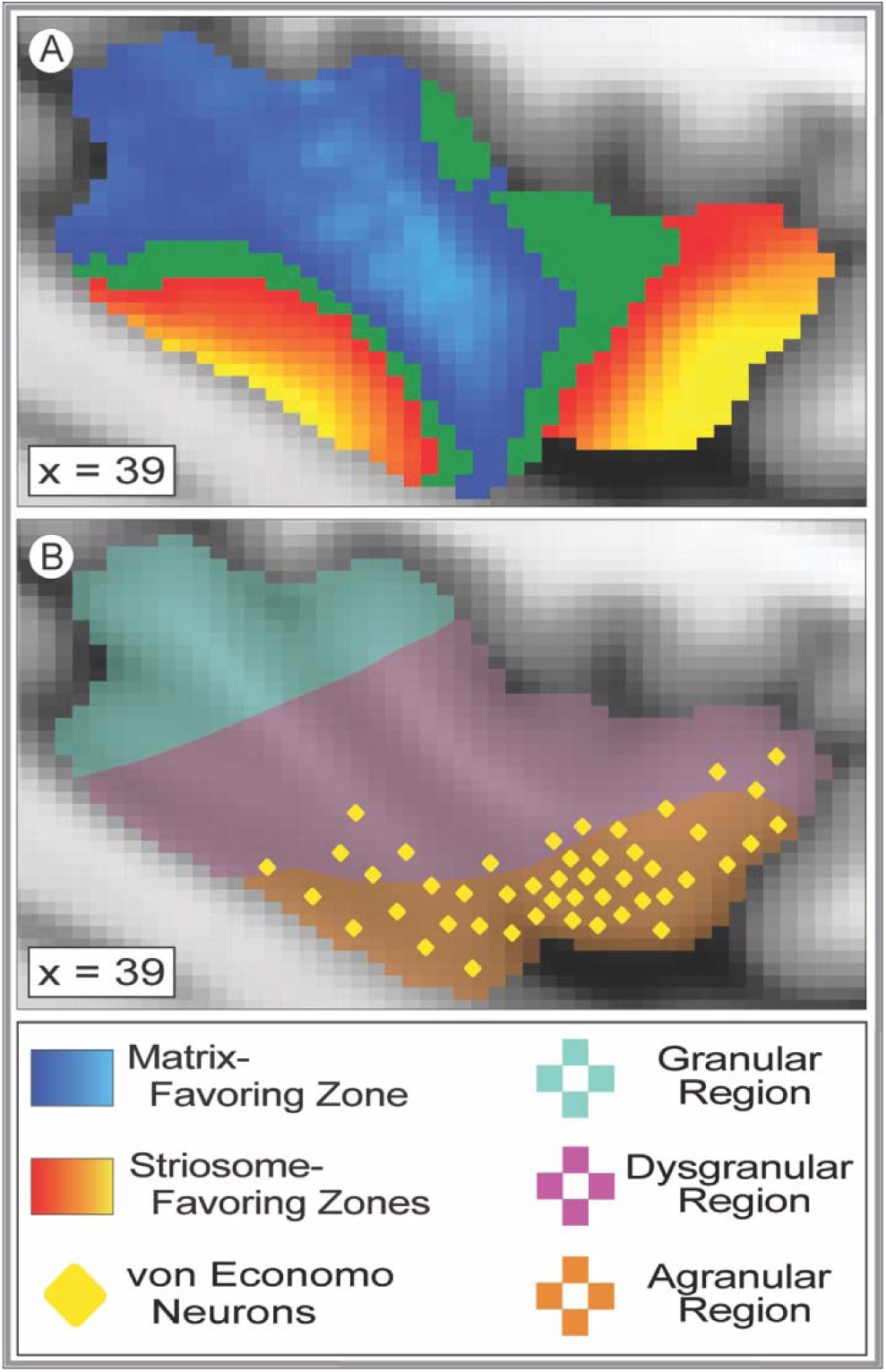
Structural connectivity with striosome-like and matrix-like voxels parallels the cytoarchitectural organization of the insula. **(A)** represents the regions in the insula with striosome-like (red-yellow) and matrix-like (blue-light blue) connectivity. The yellow and light blue colors correspond to higher degrees of connectivity (with striosome-like or matrix-like voxels, respectively). The insular mask can be seen in the background at points where connectivity was not significantly biased toward one compartment (green voxels). **(B)** represents the general cytoarchitecture of the human insula based on prior figures (Bonthius et al., 2005; ^79^ ^5,73^. The rostroventral red region represents the agranular region of the insula, the purple region represents the dysgranular region of the insula, and the blue region represents the granular region of the insula. The yellow diamonds represent the distribution of von Economo neurons in the human insula based on prior figures 5^,73,80^. The plane of visualization for both A and B is sagittal in the right hemisphere at x = 39.

### 4.5: Relating biases in structural connectivity to functional specialization

The gradient of granularity from the rostroventral to caudodorsal insula has many similarities with subregional patterns in functional connectivity. The rostroventral insula is part of a functional network that includes the frontal and temporal opercula, inferior frontal gyrus, temporal sulcus, and the anterior cingulate cortex.^82^ Recognized functions involving these regions include processing of emotions, executive functions, and cognitive control tasks.^83–85^ The caudodorsal insula is part of a functional network that includes the primary and secondary somatomotor cortices, parietal opercula, the supplementary motor area, lateral temporal cortex, and the inferior frontal gyrus.^82^ In concert with these regions, the caudodorsal insula mediates motor tasks and somatosensory processing.^3,86,87^

This rostroventral-to-caudodorsal shift in insular functional connectivity closely approximates the shift in compartment-specific biases we identified in the insula. Joel et al.^88^ argued that the matrix compartment is preferentially responsible for action, whereas the striosome compartment is preferentially responsible for critique of that action. The determination that the rostroventral insula (biased toward striosome-like voxels, Figure 3) is responsible for processing emotions and regulating behavior^4,89^ is consistent with the compartment-specific bias we observed, as the striosome receives selective projections from limbic areas, including the basolateral amygdala,^29^ rostral anterior cingulate,^25^ and prelimbic and infralimbic cortices.^57^ Additionally, prior observations that the caudodorsal insula (our matrix-favoring area) is functionally coupled with areas activated during sensory and motor tasks^82,87,90^ parallels observations in cats, squirrel monkeys, macaques, and mice that primary motor and sensory cortices selectively project to the matrix. ^50,53,54,91^ The differences in structural connectivity we identified here are potential anatomic substrates for the functional specialization of these cytoarchitecturally-distinct zones.

### 4.6: The influence of hemisphere and sex on compartment-specific projections

Cabeen et al. (2023) identified a hemisphere-by-sex interaction in insular development, influencing both cortical surface geometry and microstructure of the frontoinsular cortex (FI).^46^ They observed that women have an increased FI surface area (+7.4%) and volume (+15.3%), relative to men, in the left hemisphere. In parallel, prior fMRI and probabilistic tractography studies identified higher total insulo-striate connectivity in the left hemisphere compared with the right.^92,93^ These findings led us to investigate the interaction of hemisphere and sex on our compartment-specific structural connectivity. Similarly, to these prior reports, we identified a significant increase in striosome-favoring streamlines in the left hemisphere, specific to the rostral insula and driven by subject sex (Figure 2). It is possible that the left hemispheric dominance of insulo-striate connectivity we described may derive from the relative increase in left rostral insula volume described in women.^46^ However, this asymmetry persisted in our streamline measures after correcting for insular volume, suggesting that increased structural connectivity in the left insula is not solely a result of increased insular volume.

### 4.7: Insulo-striate dysfunction in disease and injury

We demonstrated that insular subregions selectively project to somatotopically-defined portions of striosome or matrix. Therefore, injury to either insula or striatum may produce selective functional deficits within insulo-striate networks that manifest as disease-specific symptoms, and injury may be shared among the nodes in this network. In Huntington disease (HD), comorbid striatal and insular atrophy was a potent negative predictor of performance on executive tasks relative to individuals with isolated striatal atrophy.^94^ In patients with Parkinson disease (PD) with mild cognitive impairment, dopamine depletion in the associative striatum was predictive of bilateral dopamine D2 receptor loss in the insula – suggesting that striatal and insular alterations may be interrelated in patients with these symptoms.^95^ Knowledge of compartment-specific insulo-striate connectivity patterns could provide an anatomic substrate to explain the different clinical courses of distinct HD and PD subtypes.

Linked insulo-striate functions may also inform addictive behaviors. When subjects who were avid smokers suffered from strokes that injured the insula, they were significantly more likely to quit compared with smokers with insula-sparing strokes.^96,97^ Intriguingly, the probability of smoking cessation increased when the striatum was damaged along with the insula, compared with injury to non-striatal sites. In individuals suffering from cocaine dependency, functional connectivity between the bilateral putamen and posterior insula/right postcentral gyrus is reduced.^98^ Whether this abnormal insulo-striate functional connectivity is compartment-specific is unknown. However, given that striosome and matrix have opposing responses to striatal dopamine,^21^ and the striosome, but not the matrix, innervates the dopaminergic neurons of the substantia nigra pars compacta,^99^ it is plausible that hyperdopaminergic injury and adaptation are compartment-specific. In rodent models, cocaine-induced dopamine release is significantly different in striosome and matrix, in a pattern that reverses between dorsal and ventral striatum (dorsal: 36% lower in striosome; ventral: 64% higher in striosome).^100^ Methamphetamine exposure leads to a significant depletion of monoamine terminals and neuronal apoptosis,^101^ and methamphetamine-associated injury was 4-fold greater in the striosome than the matrix.^102^ This inter-compartment difference in susceptibility was significantly larger in the medial than the lateral striatum. Since cortico-striate projections to each compartment are somatotopically organized,^56,25^ including in humans,^30^ toxin-associated neural injury may selectively impact particular insulo-striate projections, and thus, the neuropsychiatric functions of particular insular subregions. Further mapping of the somatotopy of insulo-striate projections may localize the insular functions at risk from acquired striatal injury.

### 4.8: Conclusions

Understanding the networks in which a brain region participates, and the functions those networks subserve, is critical to understanding how that region operates in health and disease. We have demonstrated that closely approximated insular subregions can have markedly different projections to the striatum, based in part on whether streamlines from those subregions target voxels with striosome-like or matrix-like connectivity profiles. These striosome-favoring and matrix-favoring insular zones may therefore be embedded in distinct CSTC loops with opposing responses to striatal dopamine release, differential susceptibility to hypoxic and excitotoxic injury, segregated vascular supplies, and that project to, and receive projections from, different sets of thalamic nuclei. A granular characterization of compartment-specific biases in structural connectivity is an essential step in understanding the functions of the human insula.

## Statements Relating to Ethics and Integrity Policies

### Data availability statement

Data used in the preparation of this manuscript was obtained from the National Institute of Mental Health Data Archive (NDA). The NDA is a collaborative informatics system created by the National Institutes of Health to provide a national resource to support and accelerate research in mental health. Dataset identifiers can be accessed through a study-specific NDA Data Object Identifier: 10.15154/1528201. This manuscript reflects the views of the authors and may not reflect the opinions or views of the NIH or of the Submitters submitting original data to NDA. Researchers can request access to de-identified MRI data by preparing an institutional Data Use Agreement for this institution. Linux code and regional brain segmentations are available for download through GitHub.

### Author Contributions

Funk, AT: The author contributed to data collection/processing, produced the first draft of the manuscript, and revised of the paper.

AAO, Hassan: The author contributed to data collection and revised the paper.

Waugh, JL: The author was the Principal Investigator, contributed to data collection and processing, authored the code used in the analyses described, and revised of the paper.

### Ethics approval statement

All research was conducted in accordance with the principles set forth in the Declaration of Helsinki. All data collection was approved by, and experimental oversight was conducted by, the Institutional Review Board for the respective institution where the subject was recruited.

### Patient consent statement

All human MRI imaging collected for this study was done with willing and informed consent and approved by the Institutional Board for the respective institution where the subject was recruited.

### Permission to reproduce material from other sources

No materials are reproduced from outside sources in this manuscript.

### Clinical trial registration

This study did not involve any clinical trial.

### Supplementary Material Access

Access to the supplementary material referenced in this study can be found at: github.com/jeff-waugh/Striatal-Connectivity-based-Parcellation.

